# Single-cell multi-omics reveals elevated plasticity and stem-cell-like blasts relevant to the poor prognosis of *KMT2A*-rearranged leukemia

**DOI:** 10.1101/2020.12.06.413930

**Authors:** Changya Chen, Wenbao Yu, Fatemeh Alikarami, Qi Qiu, Chia-hui Chen, Jennifer Flournoy, Peng Gao, Yasin Uzun, Li Fang, Yuxuan Hu, Qin Zhu, Kai Wang, Clara Libbrecht, Alex Felmeister, Isaiah Rozich, Yang-yang Ding, Stephen P. Hunger, Hao Wu, Patrick A. Brown, Erin M. Guest, David M. Barrett, Kathrin M. Bernt, Kai Tan

**Author notes:** Equal contribution (Changya Chen, Wenbao Yu). Corresponding authors Kai Tan, Kathrin Bernt, David Barret.

## Abstract

Infant ALL is a devastating malignancy caused by rearrangements of the *KMT2A* gene (*KMT2A-r*) in approximately 70% of patients. The outcome is dismal and younger age at diagnosis is associated with increased risk of relapse. To discover age-specific differences and critical drivers that mediate the poor outcome in *KMT2A-r* ALL, we subjected *KMT2A-r* leukemias and normal hematopoietic cells from patients of different ages to multi-omic single cell analysis using scRNA-Seq, scATAC-Seq and snmC-Seq2. We uncovered the following critical new insights: Leukemia cells from infants younger than 6 months have a greatly increased lineage plasticity and contain a hematopoietic stem and progenitor-like (HSPC-like) population compared to older infants. We identified an immunosuppressive signaling circuit between the HSPC-like blasts and cytotoxic lymphocytes in younger patients. Both observations offer a compelling explanation for the ability of leukemias in young infants to evade chemotherapy and immune mediated control. Our analysis also revealed pre-existing lymphomyeloid primed progenitor and myeloid blasts at initial diagnosis of B-ALL. Tracking of leukemic clones in two patients whose leukemia underwent a lineage switch documented the evolution of such clones into frank AML. These findings provide critical insights into *KMT2A-r* ALL and have potential clinical implications for targeted inhibitors or multi-target immunotherapy approaches. Beyond infant ALL, our study demonstrates the power of single cell multi-omics to detect tumor intrinsic and extrinsic factors affecting rare but critical subpopulations within a malignant population that ultimately determines patient outcome.

## Introduction

The mixed lineage leukemia 1 (*MLL1*) gene (now renamed lysine methyltransferase 2A, or *KMT2A*) encodes a nuclear protein with multiple functional domains involved in epigenetic and transcriptional regulation. Homozygous deletion of *Kmt2a* in murine embryos results in lethality at E10.5-12.5 and abnormal fetal hematopoiesis (Yagi et al., 1998; Yu et al., 1995). Rearrangements of *KMT2A* gene (*KMT2A-r*) occur with over 130 different KMT2A translocation partners (Meyer et al., 2018) and are found in acute myeloid leukemia (AML), acute lymphoblastic leukemia (ALL), mixed lineage, and biphenotypic leukemias. *KMT2A-r* leukemias account for approximately 10% of acute leukemias in all age groups (Muntean and Hess, 2012). *KMT2A* fusions are powerful oncogenes, and *KMT2A-r* leukemias typically have very few co-occurring mutations. KMT2A fusion proteins induce distinct transcriptional and epigenomic profiles (Armstrong et al., 2002; Bernt et al., 2011; Chen et al., 2015; Okada et al., 2005). Furthermore, *KMT2A-r* ALL possesses an unusual degree of lineage plasticity, which manifests as the frequent co-expression of myeloid markers as well as occasional “lineage switches”, where initially lymphoid or myeloid leukemias relapse as acute leukemias of the opposite lineage (Rossi et al., 2012).

Infant ALL represents a distinct subtype of ALL with a dismal prognosis. Approximately 70% of all infant ALL are *KMT2A-r*. The most common fusion partners in infant ALL are *AFF1 (AF4)* and *MLLT1 (ENL)*, followed by *MLLT3 (AF9)* and *MLLT10* (*AF10)*. Infant ALL is characterized by a particularly quiet genomic landscape with the predominant leukemic clone carrying a mean of 1.3 mutations (Andersson et al., 2015). The most common co-occurring mutations affect RAS and PI3K signaling, but these are often subclonal, can be lost at relapse, and do not seem to impact outcome.

Advances in risk-adapted chemotherapy, the introduction of novel targeted agents such as tyrosine kinase inhibitors for Philadelphia chromosome (Ph) positive (Ph+) ALL, and improved supportive care have increased the survival for children with ALL older than one year to more than 90% (Tasian and Hunger, 2017). In contrast, the event-free survival (EFS) for *KMT2A-r* infant ALL is 36% and has not changed in decades (Brown et al., 2019; Pieters et al., 2019). One of the strongest predictors of outcome in infant ALL is the age at diagnosis, which has been observed in multiple studies worldwide, regardless of therapeutic approach (Pieters et al., 2007, 2019). EFS rates vary dramatically between infants < 3 months of age (20-25%) and those > 6 months of age (~65%). Of note, this data includes infants with and without *KMT2A-*r and outcomes for infants with *KMT2A-*r in these age brackets are even worse.

Several strategies have been explored in recent years to improve the survival of children with *KMT2A-r* ALL. However, despite encouraging data in defined small subgroups of patients, neither the incorporation of intensive, myeloid-type chemotherapy (Interfant-06 (Pieters et al., 2019)), nor hematopoietic stem cell transplantation (Pieters et al., 2019) nor targeted therapies such as FLT3 inhibition have improved overall outcomes in large multi-center trials (Brown et al., 2016). Immunotherapies such as bispecific antibody (blinatumomab) and CAR-T cell therapies are currently revolutionizing the care of children with relapsed / refractory ALL and are emerging as strategies in the front-line therapy of very high risk disease. However, there are several challenges to developing immunotherapy strategies for infants, including the immature nature of infant T cells, inherent plasticity of *KMT2A-r* ALL, and limited amount of T cell apheresis products for CAR-T cell manufacture.

In this study, we leverage single cell multi-omic profiling to gain insights into the cellular and molecular factors that drive many of the unique features of infant ALL discussed above. In particular, we focus on understanding the developmental heterogeneity of leukemic cells, interactions between leukemic cells and immune cells, and plasticity of leukemic cells following chemotherapy and/or immunotherapy.

## Results

### Single cell multi-omics characterization of *KMT2A-r* leukemia and healthy bone marrow hematopoiesis in children

To investigate the transcriptomic and epigenetic heterogeneity in *KMT2A-r* leukemia, we performed single-cell RNA sequencing (scRNA-Seq), single-cell assay for transposase-accessible chromatin sequencing (scATAC-Seq) using 10x genomics and single-nucleus methylcytosine sequencing version 2 (snmC-Seq2) (Luo et al., 2018) on sorted leukemic blasts (CD19+) and normal hematopoietic cells (CD45+CD19-) from 25 infant and pediatric *KMT2A-r* ALL patients, including 18 infants diagnosed before one year of age (infant cohort), and 7 patients older than one year at diagnosis (pediatric cohort) (Figure 1 A-C, Table S1, STAR Methods). To confirm mutation status of malignant cells, we also performed single-cell targeted long-read sequencing (scTLR-Seq). To build a reference developmental trajectory of hematopoiesis, we sequenced hematopoietic cells from the bone marrow of five healthy pediatric donors aged from 1 to 20 years (healthy cohort, Table S1). The frequency of *KMT2A* translocation partners in our patient cohort is representative of the overall reported frequency (Meyer et al., 2018), with *AFF1* (7), *MLLT1* (5), *MLLT3* (4), and *MLLT10* (4) being the most frequent fusion partners (Figure S1A). We enriched for CD19-cells (Figure 1B, STAR Methods) to capture normal immune cells from the patients. After quality assessment and removal of low-quality cells, we sequenced a total of 200,756, 154,779, and 2,006 cells using scRNA-Seq, scATAC-Seq, and snmC-Seq2 protocols, respectively (Figure S1, STAR Methods). On average, 2,008 genes were detected per cell in the scRNA-Seq data, and 17,828 unique chromatin accessible fragments were mapped per cell in the scATAC-Seq data, 70.6% of which were mapped to peaks (Figure S1C and S1D). For snmC-Seq2 data, we sequenced an average 6.35 million reads per cell, 61.7% of which were uniquely aligned to the human genome, giving an average of 5.97% genome coverage per cell (Figure S1F). The average CpG methylation rate per cell is 80.8% (Figure S1F). These QC metrics are comparable to the published snmC-Seq2 data (Luo et al., 2017, 2018; Mulqueen et al., 2018).

**Figure 1.**
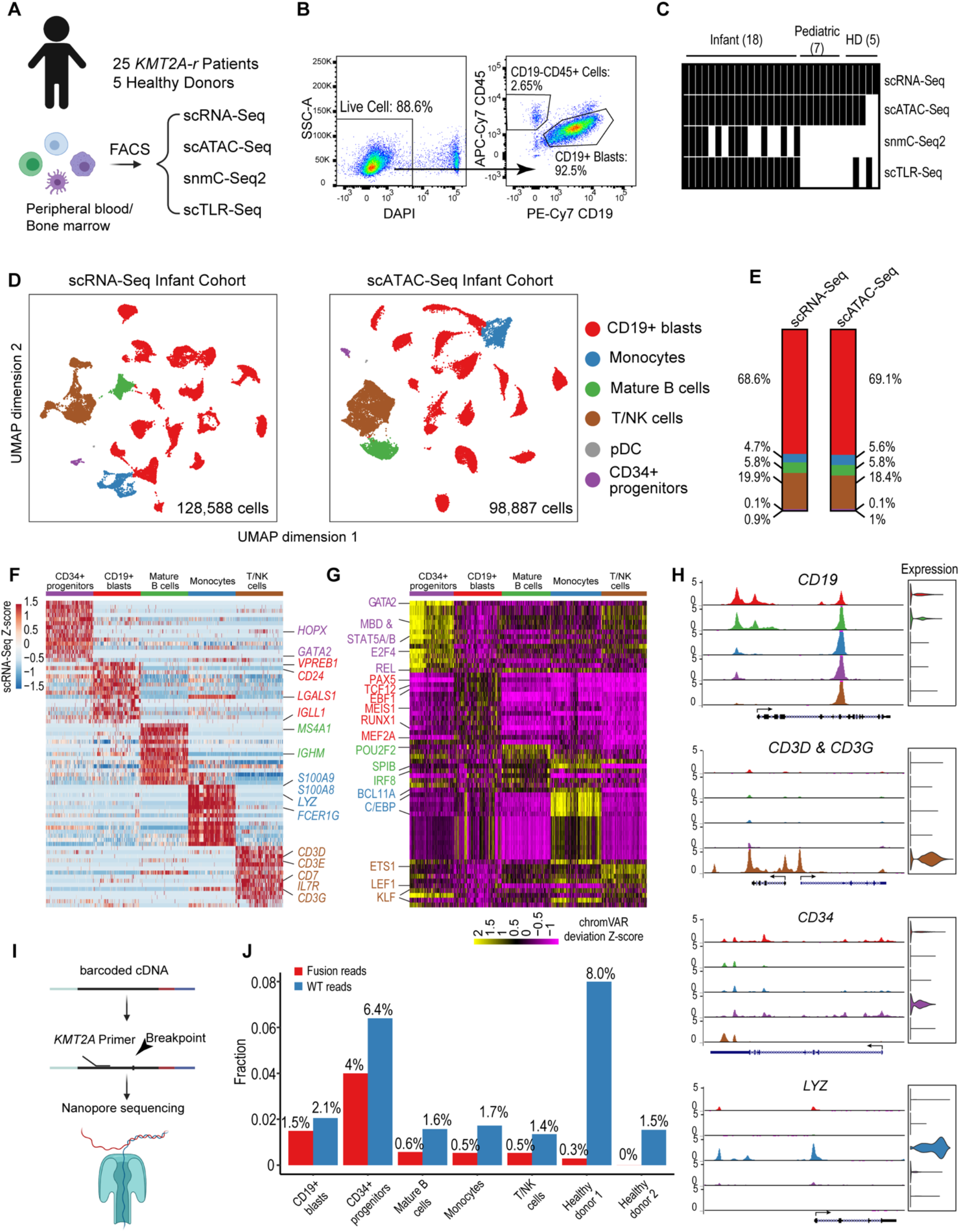
Single-cell multi-omics profiling of pediatric *KMT2A-r* leukemia. **A)** Experimental design of multi-omics profiling of *KMT2A-r* leukemia and healthy donor samples. **B)** Sorting strategy for capturing blasts and non-malignant cells from B-ALL patients. **C)** Number of assays/samples performed for each single-cell omics protocol. **D)** Overall UMAP of all scRNA-Seq cells (left panel) and all scATAC-Seq cells (right panel) of 18 infant ALL samples, colored by assigned cell populations. Total numbers of sequenced cells are indicated. **E)** Cell type compositions based on the scRNA-Seq and scATAC-Seq data in panel D. **F)** Heatmap of differentially expressed genes for each cell population compared to the rest of populations (abs(Log2(FC)) > 0.5 and FDR < 0.05). Values in the heatmap are row-wise Z-cores. Color code for each cell population is the same as in panel D. **G)** Heatmap of enriched TFs at ATAC-Seq peaks. Enrichment is represented by the normalized deviation scores (z-score) calculated by chromVAR. **H)** Genome browser tracks and gene expression violin plots for representative cell-type-specific marker genes. Left panels, aggregated scATAC-Seq signals for each assigned cell type. Right panels, normalized scRNA-Seq expression values for the corresponding cell type. **I)** Schematic of scTLR-Seq for detecting fusion transcripts in single cells. **J)** Fraction of cells with *KMT2A* fusion and wild-type reads for each cell population, including blasts, mature B cells, monocytes, NK/T cells from patients and hematopoietic cells from healthy donors. Fraction was calculated as the ratio of the number of cells with *KMT2A* reads versus the total number of sequenced cells of a given population.

Using our single-cell data, we first identified six major groups of cell types based on marker gene expression (Table S2) and visualized them using Uniform Manifold Approximation and Projection (UMAP) for both scRNA-Seq and scATAC-Seq (Figure 1D), including CD19+ blasts, T and NK cells, monocytes, mature B cells, CD34+ hematopoietic stem and progenitor cells, and plasmacytoid dendritic cells (pDCs). Further examination of marker gene expression and promoter chromatin accessibility patterns confirms the accuracy of our cell type assignment (Figure 1H). For instance, the known B-cell surface antigen, *CD19,* is specifically expressed in leukemic blasts and mature B cells. The cell type compositions are highly concordant between scRNA-Seq and scATAC-Seq data (Figure 1E).

We observed a large inter-tumor heterogeneity among blasts across the patient cohort. In contrast, normal cells (CD45+CD19-) from different patients are well mixed, suggesting that the observed heterogeneity among blasts from different patients is not due to batch effect (Figure S1E). To further confirm and identify genes that contribute to the global inter-tumor heterogeneity, we calculated the standard deviation of average gene expression in CD19+ blasts from the 18 infant patients (Figure S2A). The inter-tumor heterogeneity was indeed supported by the expression profile of known KMT2A fusion target genes or genes known to be highly expressed in *KMT2A-r* leukemia, such as *HOXA9* (Krivtsov et al., 2008; Milne et al., 2005; Muntean et al., 2009; Zeisig et al., 2004), *IRX1*, and *LGALS1* (Juszczynski et al., 2010; Paz et al., 2018) (Figure S2A). Furthermore, pathway enrichment analysis of the top 50 most variant genes revealed multiple B-cell developmental pathways and drug response pathways (Figure S2B). Among the top variant genes, *IGLL1* (*λ*5), an important marker gene for early B cell development (Herzog et al., 2009), was predominantly expressed in blasts from younger patients (< 6 mo. old, 7/11) (Figure S2C), suggesting a strong link between B-cell development and oncogenesis in this disease. Similar to expression heterogeneity, we also observed inter-tumor heterogeneity in chromatin accessibility across the patient cohort. Figure S2D shows an example of heterogeneous chromatin accessibility at an enhancer near the *LYZ* gene, a canonical myeloid lineage marker. This heterogeneity in epigenetic priming of myeloid potential is intriguing in light of the known potential of *KMT2A*-r leukemias to undergo a lineage switch to AML.

Differential gene expression and transcription factor (TF) motif chromatin accessibility analyses showed distinct transcriptional and epigenetic programs between CD19+ blasts and other cell types. Expectedly, genes that are involved in B cell development, such as *VPREB1*, *IGLL1*, and *CD24*, were expressed significantly higher in CD19+ blasts (Figure 1F). Similarly, several early B-lineage TFs were enriched at open chromatin regions specific to CD19+ blasts, such as PAX5, EBF1, TCF12 (HEB) (Figure 1G). In contrast, key lineage-specific TFs were enriched in other cell types, such as GATA2 in hematopoietic progenitors, CEBPA and CEBPB in monocytes (Zhu et al., 2016), LEF1 in T cells, and POU2F2 (OCT2) and SPIB in mature B cells (Hodson et al., 2016; Matthias and Rolink, 2005).

The blast population was first defined based on a blast gene expression signature (CD19+ blasts, Table S2). To confirm that these cells are indeed malignant, we developed the single-cell targeted long-read sequencing (scTLR-Seq) protocol which uses Oxford nanopore sequencing technology to directly sequence cDNA of fusion transcripts from individual cells (STAR Methods, Figure 1I). Our scTLR-Seq data confirmed that the blasts identified based on marker gene expression are indeed malignant cells because the fraction of fusion transcript reads in the putative blasts was significantly higher than that of normal cells from patients and bone marrow hematopoietic cells from healthy donors, which only showed a baseline fraction of fusion reads (Figure 1J).

### Leukemic blasts in younger patients have increased developmental heterogeneity

Previous studies have suggested that blasts of different B-ALL subtypes have distinct developmental origins (Castor et al., 2005; le Viseur et al., 2008). However, the landscape of perturbed developmental stages remains unknown for pediatric *KMT2A-r* leukemia. We reasoned that a normal hematopoietic developmental trajectory would be beneficial to answering this question. We therefore generated scRNA-Seq and scATAC-Seq data using bone marrow samples from five healthy pediatric donors. Given the rarity of hematopoietic progenitor and stem cells, we enriched these cells by sorting Lin-CD34+ and Lin-CD34+CD38-cells from the bone marrow samples (Figure S1B). In total, we sequenced 33,824 and 21,573 high-quality cells using scRNA-Seq and scATAC-Seq, respectively (Figure S1C & D). We then generated reference trajectories based on the transcriptome and chromatin accessibility data separately, and annotated cell populations using a manually curated marker gene list that covers the entire hematopoietic trajectory, from hematopoietic stem and progenitor cells (HSPCs) to terminally differentiated cells (Figure 2A, 2B, Table S2, STAR Methods). Both RNA- and chromatin-based trajectories identified all major hematopoietic cell types and are highly concordant (Figure 2A, Figure S3A, 3B). In particular, the B-lineage trajectory covered multiple developmental stages spanning from multipotent hematopoietic progenitors, to common lymphoid progenitors, pre-pro-B, Pro-B, Pre-B, immature B, mature B and plasma B cells.

**Figure 2.**
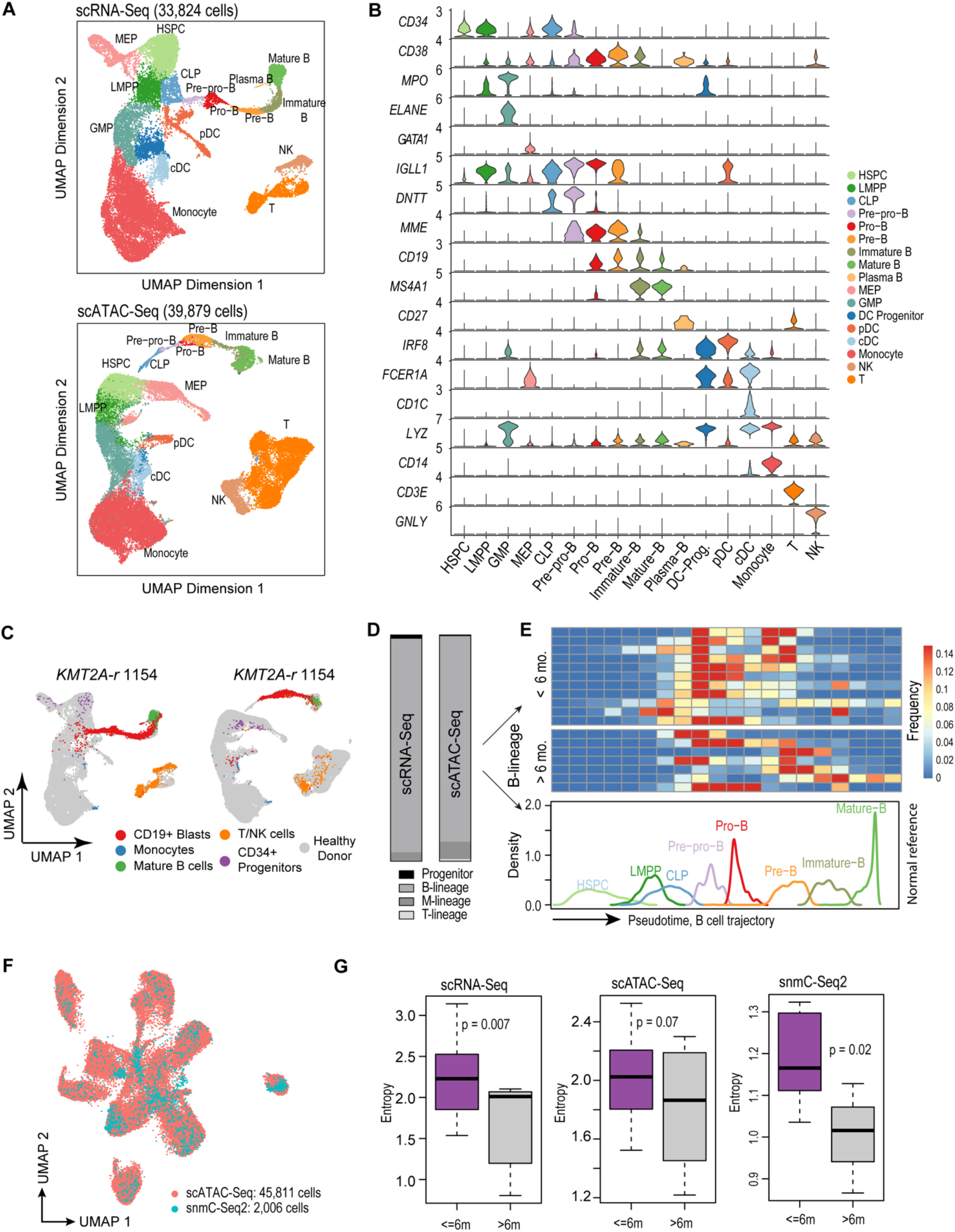
Projection of *KMT2A-r* leukemia cells to normal hematopoietic trajectory reveals larger intra-tumor heterogeneity in younger patients. **A)** UMAPs based on scRNA-Seq (top panel) and scATAC-Seq data from healthy pediatric donors (bottom panel). scATAC-Seq includes the dataset generated in this study and previously published datasets (Granja et al., 2019). Cell type annotation for scATAC-Seq data was transferred from scRNA-Seq data using Seurat. Total numbers of sequenced cells are indicated on top. **B)** Violin plots of marker gene expression used for defining the hematopoietic cell populations. **C)** Projection of patient cells onto normal hematopoietic trajectories. Left panel, representative projection of patient 1154 scRNA-Seq data. Right panel, representative projection of patient 1154 scATAC-Seq data. Grey dots, cells from healthy donors; colored dots, patient cells. **D)** Composition of CD19+ leukemic blasts in different hematopoietic lineages based on projected scRNA-Seq data (left bar) and scATAC-Seq data (right bar) data. M-lineage, myeloid lineage. **E)** Frequency of B-lineage blasts from 18 infant *KMT2A-r* patients stratified along the normal hematopoietic trajectory (pseudotime). B-lineage pseudotime from HSPCs to mature B cells is ordered into 20 bins. Upper heatmap shows the frequency of each bin from all 18 infant *KMT2A-r* patients. Lower line plot shows the frequency of each developmental stage along the pseudotime trajectory using healthy donor data. **F)** UMAP shows co-embedded snmC-Seq2 and scATAC-Seq data. Total numbers of sequenced cells of each modality are indicated. **G)** Blasts from younger patients (< 6 mo. old) show higher developmental heterogeneity based on scRNA-Seq, scATAC-Seq, and snmC-Seq2 data, respectively. Heterogeneity was quantified using Shannon’s entropy. P-values are based on *t*-test.

Next, we devised a method to project leukemic blasts from patients to the normal hematopoietic trajectory (STAR Methods). As an example, Figure 2C shows that blasts in patient 1154 were mostly arrested after the pre-pro B stage with a minor fraction of blasts arrested at the earlier multipotent hematopoietic progenitor stage. The projections based on the transcriptome and chromatin accessibility data are highly concordant (Figure S3C). Overall, >95% leukemic blasts were projected onto the B-cell developmental trajectory across the 18 patients (Figure 2D). However, within this trajectory, there is a wide spectrum of developmental stages at which blasts from the same patient were arrested, from HSPCs to immature B cells (Figure 2E). We quantified this intra-tumor developmental heterogeneity using Shannon’s entropy. Remarkably, we found that blasts from younger patients (< 6 months old) exhibited significantly higher developmental heterogeneity that manifested at both the transcriptomic (p = 0.007) and chromatin accessibility (p = 0.07) levels (Figure 2G). We next integrated our snmC-Seq2 data with scATAC-Seq data (Figure 2E, STAR Methods). Using the DNA methylation data, we again observed higher developmental heterogeneity in the blasts of younger patients (p = 0.02) (Figure 2G, right panel).

### Transcriptomic and epigenomic signatures of blasts in younger patients

Given the significantly higher lineage plasticity among blasts in younger patients, we next sought to identify genes and epigenomic features underlying this difference. Using the cutoff of abs(log_2_FC) > 0.5 and FDR < 0.05, we identified 495 genes that were differentially expressed (DEGs) in blasts between younger and older patients in at least one B-cell developmental stage (Figure 3A, Table S5). Among those DEGs, 360 were expressed higher and 213 were expressed lower in younger patients, respectively. Clustering analysis of the DEGs revealed five clusters. Genes in cluster 1 were expressed higher in younger patients across the entire B-cell developmental trajectory. Among the top enriched pathways are response to viral infection, antigen processing and presentation and AP-1 pathway (Figure 3C, left panel). Other genes in cluster 1 include several key B-cell development genes, such as *IGLL1*, *VPREB1*, *IL7R*. Genes in cluster 4 were expressed higher in younger patients and predominantly at immature B stage. The top enriched pathways include ribosome biogenesis, metabolism and translation (Figure S4A). Genes in clusters 5 and 2 were expressed lower in younger patients in early (progenitor and pre-pro-B stages) and late (pre-B and immature B) B-cell developmental stages, respectively. The top enriched pathways in cluster 2 and cluster 5 are hematopoietic cell lineage (Figure S4A) and response to corticosteroid (Figure 3C, right panel), respectively.

**Figure 3.**
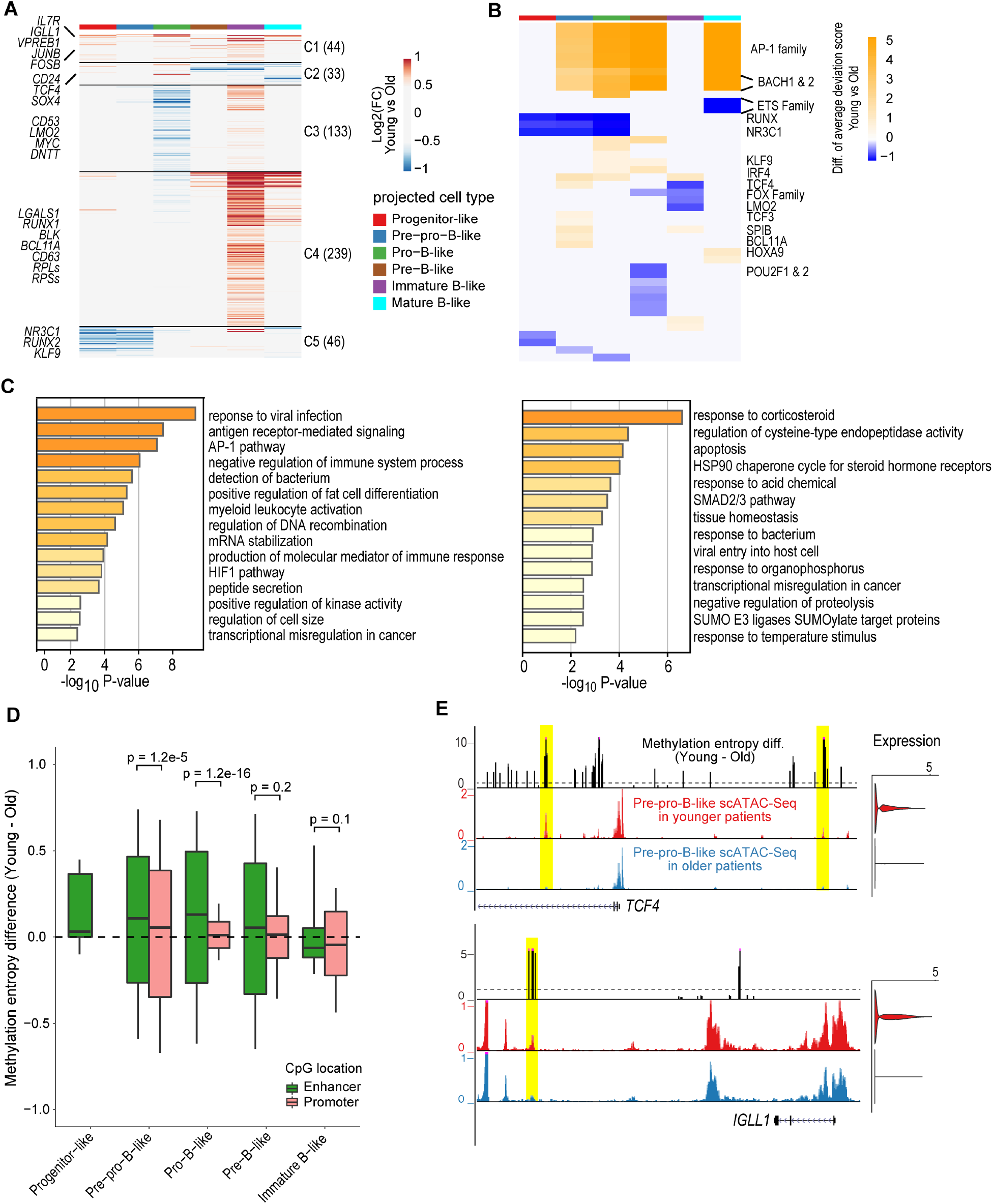
Transcriptomic and epigenomic signatures of blasts in younger patients. **A)** Heatmap for differentially expressed genes (abs(log_2_FC) > 0.5 & FDR < 0.05) of blasts arrested at various B-cell developmental stages between younger (< 6 mo. old) and older patients (> 6 mo. old). DEGs were pooled and clustered by k-means clustering (k = 5) based on their log_2_FC. Number of genes in each cluster is indicated in the parenthesis. Non-significant genes are colored grey. **B)** Heatmap for differential TF motif accessibility of blasts arrested at various B-cell developmental stages between younger and older patients. For each TF in each cell, the motif accessibility at scATAC-Seq peaks were computed as the normalized deviation score using chromVAR. Color in the heatmap indicates the difference in normalized chromVAR deviation scores averaged across all cells in younger versus older patients. TFs with differential accessibility between younger and older patients were identified by Wilcoxon test of the normalized deviation scores between the two groups with a FDR cutoff < 0.05. Non-significant TFs were colored in grey. **C)** Pathway enrichment analysis results for DEGs in clusters 1 (left panel) and cluster 5 (right panel). **D)** Methylation entropy difference of CpG sites in promoters and enhancers of blasts between younger and older patients. Only promoters of DEGs and enhancers with at least one hit of the enriched TF motifs in panel B were used. P-values were based on *t*-test. There was no CpG overlapping the promoters of the DEGs in progenitor-like blasts. Mature B-like blasts were excluded from this analysis because only a few cells were found. **E)** Representative enhancers of *TCF4* and *IGLL1* with high DNA methylation entropy difference between younger and older patients. Top track, Exp(methylation entropy difference). The dashed black line indicates a value of 1 (i.e. higher entropy in younger patients). Middle and bottom tracks, aggregated scATAC-Seq signals for younger and older patients, respectively. Expression of the two genes is shown on the right.

TF motif accessibility analysis revealed 45 TFs that had genome-wide chromatin accessibility difference (Wilcoxon test FDR < 0.05) in blasts between younger and older patients in at least one B-cell developmental stage. Among those TFs, 26 had higher and 20 had lower accessibility in younger patients (Figure 3B, Table S6). Consistent with their expression levels, the genome-wide motif chromatin accessibility for the AP-1 factors were higher in younger patients. In contrast, chromatin accessibility of the NR3C1 motif was lower in younger patients.

We next examined the DNA methylation difference in blasts between younger and older patients. On average, we identified 22,432 differentially methylated regions (DMRs) for each B-cell developmental stage (Table S7). Together with the result of chromatin accessibility analysis, these results suggest that there is a large difference in the epigenome of blasts between patients of the two age groups. The DNA methylation change in gene promoter was consistent with the change in gene expression. For instance, promoters of *IL7R*, *IGLL1*, and *TCF4* were hypomethylated in younger patients at multiple B-cell developmental stages (Figure S4B), which is consistent with their increased expression. In contrast, promoters of *NR3C1*, *KLF9*, and *RUNX2* were hypermethylated in younger patients at early stages of B-cell development, with decreased gene expression at these stages (Figure S4B & C, Figure 3A & B).

Disordered methylation pattern at adjacent CpG sites has been reported to contribute to intra-tumor epigenetic heterogeneity and be associated with adverse clinical outcome (Landau et al., 2014; Li et al., 2016). We quantified this methylation disorder using DNA methylation entropy (Xie et al., 2011) (STAR Methods). Strikingly, we found that methylation entropy is higher across the genome of blasts from younger patients at multiple B-cell developmental stages (p < 0.01, Figure S4D). This pattern was also observed at cis-regulatory sequences, such as promoters and enhancers (Figure 3D). Furthermore, at earlier developmental stages, including progenitor, pre-pro-B and pro-B stages, we found that enhancers in blasts from younger patients had even higher increase in methylation entropy compared to promoters (Figure 3D), suggesting that disordered DNA methylation at enhancers may contribute more to the intra-tumor heterogeneity and clinical outcome (Figure 3D). Figure 3E shows two representative enhancers at the *TCF4* and *IGLL1* loci in pre-pro-B blasts. The enhancers had higher methylation entropy and higher chromatin accessibility in younger patients. Expression of the two putative target genes was also higher in younger patients. Taken together, our single cell DNA methylation data is concordant with our transcriptome and chromatin accessibility data and provides further evidence supporting the higher intra-tumor heterogeneity in younger patients.

Steroids such as prednisone and dexamethasone are one of the most important agents for the treatment of *KMT2A-r* ALL patients (Pieters et al., 2007). However, steroid resistance largely contributes to the poor prognosis of infant *KMT2A-r* ALL, especially among younger patients at diagnosis (Dördelmann et al., 1999; Pieters et al., 1998; Ramakers-van Woerden et al., 2004; Spijkers-Hagelstein et al., 2014). The top enriched pathway among genes in cluster 5 is response to corticosteroid and includes genes such as *NR3C1* and *KLF9* (Figure 3A). The promoters of these two genes were hypermethylated in younger patients (Figure S4C). *NR3C1* encodes the glucocorticoid receptor and previous studies have reported that *NR3C1* is important for glucocorticoid treatment response in various types of acute leukemia (Haowen et al., 2018; Wandler et al., 2020). *KLF9* has been reported as a gene that responds to glucocorticoid signals (Gans et al., 2020; Reddy et al., 2009). It is still unknown how *NR3C1* mediates glucocorticoid treatment response in different age groups of infant *KMT2A-r* ALL. Lower *NR3C1* and *KLF9* gene expression and motif accessibility in younger patients provide a potential mechanistic explanation for inferior glucocorticoid treatment response and for therapy resistance in younger patients. Our analysis further revealed higher expression of *AP-1* transcription factors and motif accessibility in younger patients. Because AP-1 factors are the target that glucocorticoids antagonize (González et al., 2000), together these findings uncovered a potential role of the AP-1 regulon in the glucocorticoid resistance in younger patients.

### A rare HSPC-like blast population exists in younger patients

We found a small population of cells that resemble hematopoietic stem and progenitor cells and represent 1.27% and 1.47% of total cells in our scRNA-Seq and scATAC-Seq data, respectively (Figure 4A). These cells are CD34+CD19-(Figure 1H) and express several canonical HSPC TFs such as *GATA2* and *HMGA2*. In contrast to CD19+ blasts, they do not express key B-cell developmental genes, such as *EBF1*, *PAX5*, *VPREB1*, and *DNTT* (Figure 4B left panel, Figure S5A). Furthermore, several key HSPC TFs were enriched but early B-cell TFs were depleted at open chromatin regions in this population compared to CD19+ blasts (Figure 4B, right panel). Finally, scTLR-Seq detected *KMT2A* fusion transcripts in this population at a similar level as CD19+ leukemic blasts (Figure 1K, Figure 4C). Taken together, multiple lines of evidence suggest that these abnormal cells are leukemic cells with hematopoietic stem and progenitor transcriptomic and epigenomic signatures. We hereafter term this population HSPC-like blast.

**Figure 4.**
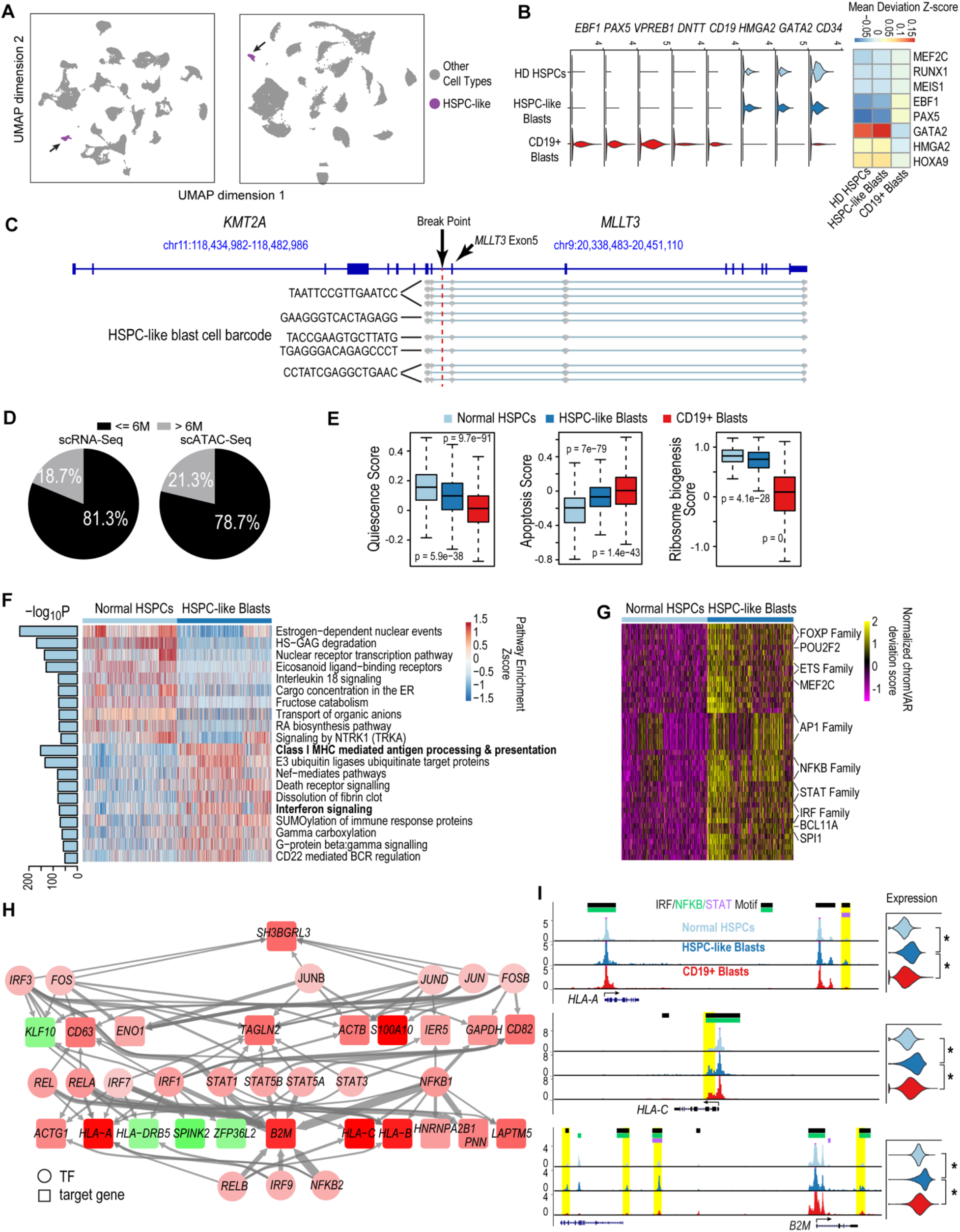
A HSPC-like blast population exists in younger patients. **A)** UMAPs of single-cell data from 18 infant patients highlighting the hematopoietic stem and progenitor-like population in the peripheral blood of the patients (HSPC-like blasts, purple). Grey, other cell types. Left panel, UMAP based on scRNA-Seq data, showing 128,588 total cells, 1136 of which are HSPC-like blasts. Right panel, UMAP based on scATAC-Seq data, showing 98,887 total cells, 1020 of which are HSPC-like blasts. **B)** Gene expression and regulator activity signatures for normal HSPCs, HSPC-like blasts and CD19+ blasts. Left panel, violin plots of marker gene expression in the three cell types. Right panel, activity of cell-type-specific transcriptional factors in the three cell types. Activity was measured as the mean TF motif chromatin accessibility score in each cell type. **C)** Representative result of fusion reads identified in HSPC-like blasts from patient 1154 using scTLR-Seq. Fusion reads from multiple HSPC-like blasts (with different cell barcodes) are shown. **D)** Percentages of total HSPC-like blasts contributed by younger and older patients, based on scRNA-Seq and scATAC-Seq data. **E)** Quiescence, apoptosis, and ribosome biogenesis signature gene scores for bone marrow HSPCs from normal donors, HSPC-like blasts, and CD19+ blasts. Scores were calculated as the sum of z-scores (across all cells) of all genes of a signature. P-values were computed using the Wilcoxon test. **F)** Pathways enriched comparing HSPC-like blasts and bone marrow HSPCs from normal donors. Enriched pathways were identified using the AUCell method. Heatmap shows the top 20 enriched pathways for HSPC-like blasts and normal HSPCs, respectively. Barplot on the left shows the adjusted p-value for enrichment. **G)** Heatmap of differential TF motif accessibility in HSPC-like blasts compared to normal bone marrow HSPCs. Values are z-score normalized deviation score calculated using chromVAR. TFs with differential accessibility between HSPC-like blasts and normal HSPCs were identified using Wilcoxon test of the normalized deviation scores between the two groups with a FDR cutoff < 0.05. **H)** Transcriptional regulation network distinguishing HSPC-like blasts from normal HSPCs. For clarity only the part of the TRN involving AP-1, NFKB, STAT, and IRF factors is shown. Nodes represent either enriched TFs or differentially expressed genes in the HSPC-like blasts versus normal HSPCs. Node color is proportional to the −log_10_(FDR) of differential expression with red being up-regulation and green being down-regulation. Edge line thickness is proportional to the −log_10_(FDR) of the linear regression coefficient for the predicted enhancer-promoter interaction. **I)** Example MHC class I genes up-regulated in HSPC-like blasts. Left panels, aggregated scATAC-Seq signals in normal HSPCs, HSPC-like blasts, and CD19+ blasts. Putative enhancers for each gene are highlighted in yellow. Motifs of STAT or NFKB or IRF transcription factors overlapping the enhancers are indicated at the top. Right panels, violin plots for normalized expression of the selected genes in the three cell types. *, adjusted p-value < 1e-10.

Intriguingly, we found that ~80% of HSPC-like blasts came from patients younger than 6 months (Figure 4D), suggesting a potential link between the presence of HSPC-like blasts and poorer prognosis in younger patients. To further characterize this population, we computed several gene signature scores based on known hallmarks of normal HSPCs, including quiescence score, apoptosis score (Kondo et al., 2003), and ribosome biogenesis score (Cai et al., 2015). Normal HSPCs are quiescent, more resistant to apoptosis and have a higher activity of ribosome biogenesis. We found that HSPC-like blasts resemble a cellular state between normal HSPCs and CD19+ blasts in terms of these three gene signatures (Figure 4E). We next performed differential gene expression analysis and identified 55 up-regulated and 17 down-regulated genes in HPSC-like blasts compared to normal HSPCs (abs log_2_FC > 0.5 and FDR < 0.05, Figure S5B). Pathway enrichment analysis revealed a number of interesting pathways that exhibited differential activities in HSPC-like blasts (Figure 4F). In particular, type II interferon response and MHC class I genes were upregulated in HSPC-like blasts. Consistent with the upregulated expression of interferon response genes, scATAC-Seq data showed the enrichment of open chromatin regions containing motifs of interferon effector TFs in HSPC-like blasts, including STAT, IRF, and NFKB (Figure 4G). Taken together, these data suggest a significant upregulation of interferon-γ response and MHC class I genes in the HSPC-like blasts. The other group of enriched TFs in HSPC-like blasts compared to normal HSPCs include critical factors for B-cell development and maturation, such as MEF2C (Herglotz et al., 2016; Kong et al., 2016), ETS factors (DeKoter et al., 2002; Eyquem et al., 2004) and POU2F2 (Hodson et al., 2016; Matthias and Rolink, 2005) (Figure 4G). Increased activity of these factors further suggests that HSPC-like blasts have a different gene regulatory program compared to normal HSPCs.

To further study the transcriptional program specifically operating in the HSPC-like blasts, we constructed a transcriptional regulation network (TRN) by integrating our scRNA-Seq and scATAC-Seq data and focusing on differentially enriched TFs and differentially expressed genes in HSPC-like blasts compared to normal HSPCs (see STAR Methods). For clarity, here we present the subnetwork involving interferon effector TFs, including AP-1, NFKB, STAT, and IRF factors (Figure 4H). The full TRN is provided in Table S8. Our subnetwork model suggests that interferon effector TFs act in a highly combinatorial fashion to regulate the MHC class I genes and other target genes in the HSPC-like blasts. For instance, we identified binding sites of multiple interferon effector TFs in enhancers of the *HLA-A*, *HLA-C* and *B2M* genes (Figure 4I).

### An immunosuppressive circuit between HSPC-like blasts and cytotoxic lymphocytes

Interferon-γ (IFNG) signaling plays an important role in tumor cells to evade the attack by cytotoxic lymphocytes, namely CD8+ T cells and Natural Killer (NK) cells. To further investigate the role of IFNG signaling between HSPC-like blasts, CD19+ blasts, and immune cells, we performed a joint analysis of patient blasts and normal immune cells. After filtering out low-quality cells, we obtained 36,313 and 29,528 normal immune cells based on scRNA-Seq and scATAC-Seq data, respectively (Figure 5A, Figure S6A). The composition of immune cell types in our patient cohort is concordant with previous observations, such as higher fraction of naive T cells in younger patients and higher fraction of effector and cytotoxic T cells in older patients (Figure S6B) (Kumar et al., 2018a; Woodland and Blackman, 2006). Natural killer T (NK T) cells and natural killer cells are the major sources of IFNG in the human body (Arase et al., 1996; Borden et al., 2007; Schroder et al., 2004). We indeed found that NK T cells (and to a lesser extent NK cells) were the major source of IFNG in our patient cohort (Figure 5B). Interestingly, we found that a higher fraction of NK T cells were identified in younger patients (Figure S6B). Among detected NK T cells, a higher fraction of them expressed *IFNG* in younger patients compared to older patients (p = 7e-5, Figure 5C). HSPC-like blasts also expressed a higher level of IFNG receptor 2 gene (*IFNGR2*) compared to normal HSPCs (p = 9e-3, Figure 5C, right panel). Taken together, these results suggest an aberrantly elevated IFNG signaling between HSPC-like blasts and NK T cells in younger patients.

**Figure 5.**
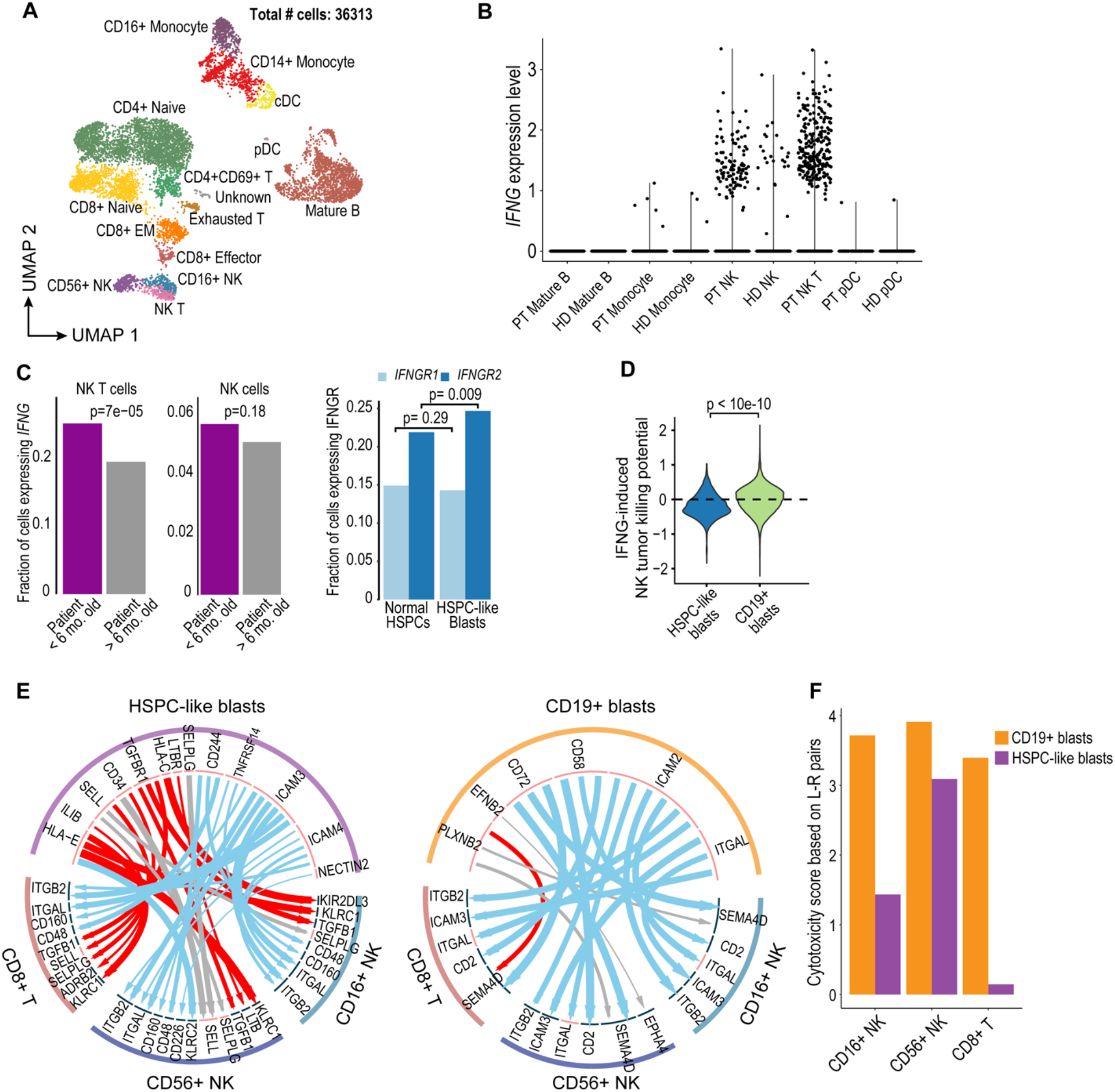
An immunosuppressive circuit between HSPC-like blasts and cytotoxic lymphocytes. **A)** UMAP of scRNA-Seq data for normal immune cells in 18 *KMT2A-r* patients. Number of sequenced cells is indicated. **B)** NK T cells are the major source of IFNG in patients. Shown are violin plots of *IFNG* expression in various immune cell populations in both *KMT2A-r* patients and healthy donors. HD, healthy donor; PT, patient. **C)** Upregulated IFNG signaling in HSPC-like blast in younger patients. Left, barplot showing younger patients have a higher fraction of NK T cells expressing *IFNG*. Right, barplot showing IFGN receptor 2 gene (*IFNGR2*) was expressed higher in HPSC-like blasts compared to normal HSPCs. **D)** Activating and suppressive signaling pathways for NK cell cytotoxicity induced by IFNG signaling in blasts. Y-axis, potential for inducing NK cell cytotoxicity based on combined normalized expression of genes in activating and suppressive pathways in HSPC-like blasts and CD19+ blasts. **E)** Predicted ligand-receptor interactions between HSPC-like blasts (left panel) or CD19+ blasts (right panel) and two major classes of cytotoxic lymphocytes, CD8+ T cells and NK cells. Red, blue, and grey arcs, suppressive, activating and unknown interactions, respectively. **F)** Cytotoxicity scores of NK and CD8+ T cells based on combined activating and suppressive signaling between the cytotoxic lymphocyte and the blast via ligand-receptor (L-R) pairs. L-R pairs were based on those in panel E. The score is the sum of mean normalized ligand and receptor gene expression for each pair of cell types over all predicted L-R pairs. Activating L-R pairs gave a positive value whereas suppressive L-R pairs gave a negative value.

IFNG signaling in tumor cells affects two important classes of genes relevant to lymphocytes. Induction of the MHC class I genes in tumor cells can function as suppressive signals for NK cell cytotoxicity whereas induction of cell adhesion genes such as *ICAM1* function as activating signals for NK cell cytotoxicity (Aquino-López et al., 2017; Raulet and Guerra, 2009). To assess the overall effect on NK cell activation due to signaling between leukemic blasts and NK cells, we computed a combined score for the induction of NK cell cytotoxicity based on the expression levels of the two classes of genes in the blasts. We found that signaling to cytotoxic lymphocytes by HSPC-like blasts has an overall immunosuppressive effect towards NK cells compared to CD19+ blasts (p < 1e-10, Figure 5D).

To further understand signaling between leukemia blasts and cytotoxic lymphocytes (CD8+ T and NK cells), we next conducted a systematic search of active ligand-receptor (L-R) pairs mediating signal transduction between these two classes of cells (STAR Methods). We performed this analysis for HSPC-like blasts and CD19+ blasts separately, using manually curated, high-quality L-R pairs (STAR Methods). Figure 5E shows the predicted L-R pairs between the two types of blasts and the two types of cytotoxic lymphocytes. Most strikingly, the majority of L-R pairs between HSPC-like blasts and the lymphocytes are known to have a suppressive role on the activity of the cytotoxic lymphocytes (Frey; Huntington et al., 2020; Martinez-Lostao et al., 2015; Russick et al., 2020). Example L-R pairs include killer Ig-like receptors (KIRs) and MHC class I proteins (Anfossi et al., 2006; Chiossone et al., 2018; Dorfman and Raulet, 1996), transforming growth factor beta receptor (TGFBR) and TGFB (Batlle and Massagué, 2019; Pickup et al., 2013; Wrzesinski et al., 2007). We next computed an overall lymphocyte cytotoxicity score (or activation score) by summing the normalized expression levels of all predicted L-R pairs between a given blast type and lymphocyte type. We found that the cytotoxic scores are lower for all three types of lymphocytes when considering signaling from HSPC-like blasts, compared to signaling from CD19+ blasts (Figure 5F). In other words, HSPC-like blasts appear to exert an overall immunosuppressive effect on these cytotoxic lymphocytes, which may facilitate their immune escape.

### Pre-existing lymphomyeloid primed progenitor and myeloid blasts in treatment-naive B-ALL predict lineage switch

Lineage switch between lymphoid and myeloid fate post therapy or relapse is rare but usually associated with poor outcome. Previous studies have shown that lineage switch predominantly occurs in *KMT2A-r* leukemia and in younger patients (Rayes et al., 2016; Rossi et al., 2012). Among our cohort of 18 treatment-naive *KMT2A-r* patients who were clinically diagnosed as B-cell precursor (BCP)-ALL, we found several of them have substantial high levels of myeloid-lineage blasts (Figure 6A) at diagnosis. For instance, patient PAYZLC had 41% and 71% myeloid-like blasts as measured by scRNA-Seq and scATAC-Seq data, respectively (Figure 6A, 6B). Overall, higher percentages of myeloid-like blasts were observed in patients based on ATAC-Seq data compared to RNA-Seq data (Figure 6A), suggesting epigenomic profiling may be a more sensitive means to detect lineage switch potential.

**Figure 6.**
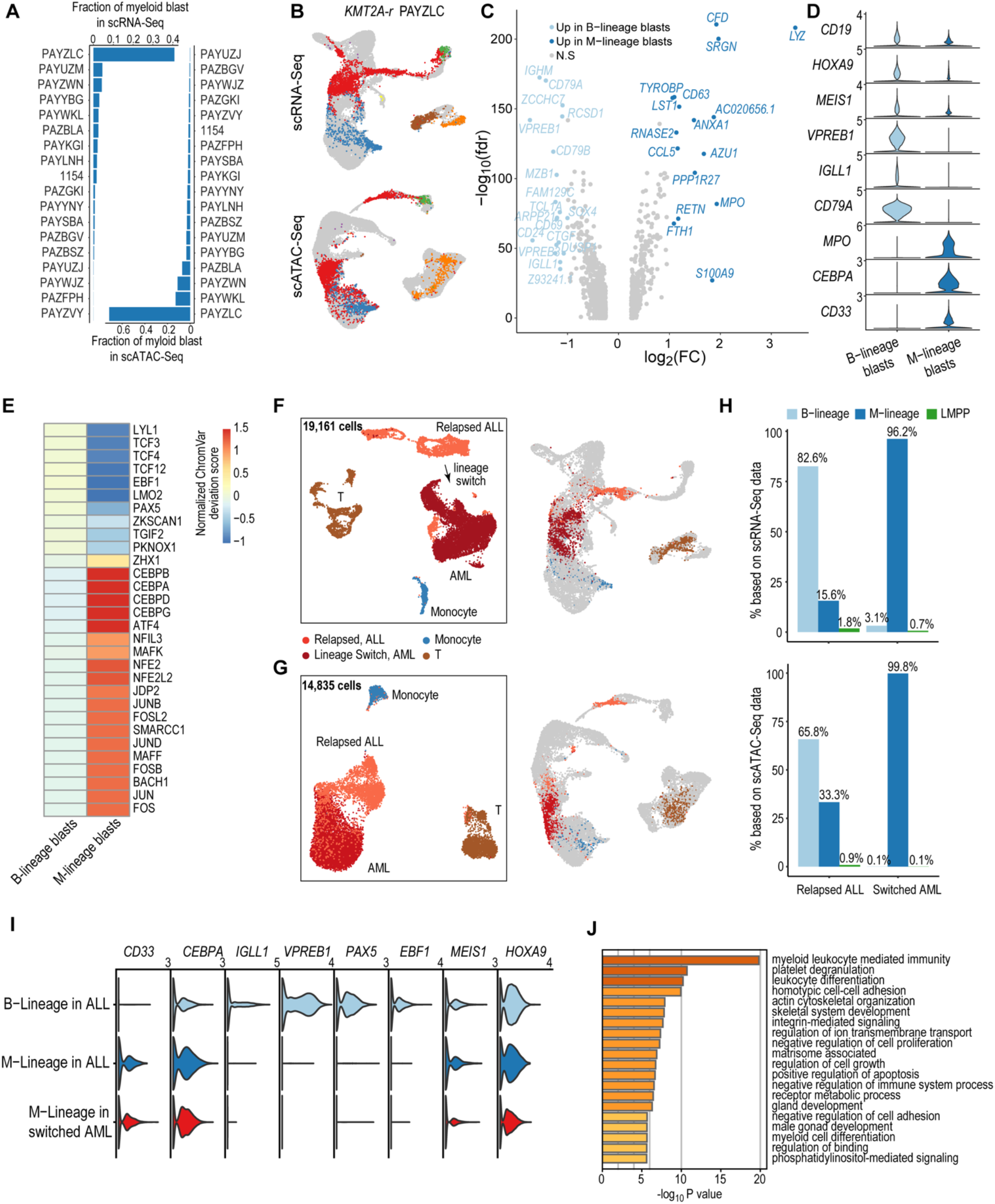
Pre-existing lymphomyeloid primed progenitor and myeloid blasts in treatment-naive patients as indicators of lineage switch. **A)** Frequencies of CD19+ blasts that were projected to the myeloid lineage (GMPs, monocytes, and dendritic cells) in all 18 infant *KMT2A-r* patients based on scRNA-Seq and scATAC-Seq data. **B)** Projection of patient PAYZLC data onto normal hematopoietic trajectory. Top panel, projection of scRNA-Seq data. Bottom panel, projection of scATAC-Seq data. Grey dots, cells from healthy donors; colored dots, patient cells. **C)** Volcano plot for differentially expressed genes between M-lineage blasts and B-lineage blasts. Analysis was based on projected blasts from all 18 patients. DEGs were identified with the cutoff of abs(log_2_FC) > 0.5 & FDR < 0.01. Those with abs(log_2_FC) > 1 are highlighted on blue. **D)** Violin plots for representative signature genes in M-lineage blasts and B-lineage blasts. **E)** Heatmap of differential TF motif accessibility in B-lineage and myeloid-lineage blasts. Analysis was based on projected blasts from 18 patients. Values are z-score normalized deviation score calculated using chromVAR. TFs with differential accessibility between B-lineage and myeloid-lineage blasts were identified using Wilcoxon test of the normalized deviation scores between the two groups with a FDR cutoff < 0.05. **F)** & **G)** UMAP of scRNA-Seq **(F)** and scATAC-Seq **(G)** data for a pediatric *KMT2A-r* patient with paired samples before and after lineage switch. Left panel, UMAP of paired samples, colored by assigned cell populations. Total numbers of sequenced cells are indicated. Right panel, projection of patient cells to the normal hematopoietic trajectory. Grey dots, cells from healthy donors; colored dots, patient cells. **H)** Fraction of B- and myeloid-lineage blasts before and after lineage switch. Top panel, fraction based on scRNA-Seq data; Bottom panel, fraction based on scATAC-Seq data. **I)** Violin plots of gene expression of B-lineage and myeloid-lineage marker genes before and after lineage switch. **J)** Enriched pathways among differentially expressed genes between normal LMPP from healthy donors and LMPP-like blasts in patient samples before lineage switch.

Compared to B-lineage blasts, these myeloid-lineage blasts differentially expressed 231 genes, many of which are known marker genes for myeloid cells, such as *MPO*, *CEBPA*, and *CD33* (Figure 6C, 6D). scATAC-Seq analysis further confirmed activity of lineage-specific TFs in the myeloid-lineage blasts (e.g. C/EBP factors and BACH1 and 2) and B-lineage blasts (e.g. EBF1 and PAX5), respectively (Figure 6E, Figure S7A). Collectively, these data strongly suggest that myeloid-lineage blasts exist in some patients at initial diagnosis of B-ALL. Outcome data for the infant cohort is not yet available. To further investigate this hypothesis, we analyzed longitudinal samples from two BCP-ALL *KMT2A-r* patients whose disease underwent a lineage switch, and for whom we had serial samples available. Patient one developed a lineage switch to AML after treatment with CART-19 for relapsed B-ALL (Figure S7B). Patient two was treated with CART-19 for relapsed *KMT2A-r* ALL, then relapsed with CD19-negative ALL, received inotuzumab (anti-CD22 monoclonal antibody) which failed to induce a deep remission, and then went on to receive CART-22 since CD22+ B-ALL persisted by flow cytometry (Figure S7C). At the next relapse, this patient’s leukemia had undergone a phenotypic lineage switch to AML. In both patients, we identified pre-existing myeloid blasts before lineage switch in our scRNA-Seq and scATAC-Seq data (Figure 6F-H; Figure S7D-F). For both patients, myeloid signature gene expression was evident in both pre-switch myeloid blasts and post-switch AML blasts (Figure 6I; Figure S7G). Interestingly, at several time points the myeloid potential by ATAC-Seq exceeded the expression of myeloid signature genes, underscoring the much greater power of single-cell epigenomic profiling to detect lineage potential compared to transcriptomic analysis.

Besides pre-existing myeloid-lineage blasts, altered differentiation of lymphomyeloid-primed progenitors (LMPP) could also contribute to lineage switch. Indeed, we identified a small fraction of LMPP-like blasts in multiple patients of our infant cohort (Figure S8A-B). These LMPP-like blasts were present both pre- and post-switch in our longitudinal samples (Figure 6H; Figure S7D). Interestingly, the fractions of LMPP-like blasts and myeloid-lineage blasts were significantly correlated across our patient cohort (Spearman correlation = 0.61 and 0.68 and p-value = 0.004 and 0.001, Figure S8C). To test the hypothesis of an altered differentiation program in LMPP-like blasts, we performed differential expression analysis. We identified 483 and 752 DEGs between normal LMPPs and LMPP-like blasts from pre- and post-lineage switch samples, respectively. Strikingly, the top enriched pathway among the DEGs in both comparisons was myeloid leukocyte mediated immunity (Figure 6J; Figure S8D). In summary, these data suggest that altered differentiation of LMPP with a myeloid-lineage bias may contribute to lineage switch under therapeutic pressure.

## Discussion

Although our patient cohort was clinically diagnosed as B-cell precursor leukemia, our single-cell multi-omics analysis revealed a large and previously unappreciated developmental plasticity of the arrested leukemic blasts, spanning the entire B-cell developmental trajectory from HSPCs to immature B cells. By performing a developmental-stage-wise comparison between arrested blasts and corresponding normal cells, we identified 148 genes that are differentially expressed across at least four developmental stages (Figure S9A-B). These genes represent a core *KMT2A-r* ALL gene signature at the single-cell resolution. Interestingly, we found that this gene list is enriched for known drug targets (Figure S9C) and therefore warrants future investigation to identify novel therapeutic targets specifically for this disease. Interesting examples include *LGALS1, DDIT4, and CTGF*. Several studies demonstrated that LGALS1 is a valuable drug target for *KMT2A-r* ALL (Juszczynski et al., 2010; Paz et al., 2018). Up-regulation of *DDIT4* is associated with poor prognosis of AML (Cheng et al., 2020; Pinto et al., 2016, 2017). *CTGF/CCN2* encodes a connective tissue growth factor and is highly expressed in several pediatric and adult ALL and is associated with poor clinical outcome (Boag et al., 2007; Sala-Torra et al., 2007).

Unexpectedly, we found blasts from patients younger than six months had significantly higher developmental heterogeneity than blasts from older patients. This observation provides a potential cellular and molecular explanation for the worse prognosis for younger patients.

Another key finding of our study is the discovery of the CD34+CD19-HSPC-like population that is observed in younger patients. Compared to normal HSPCs, this population exhibited abnormal transcriptomic and epigenomic features, such as premature upregulation of the *IGHM* gene and increased activities of several early B-cell development transcription factors (Figure 4, Figure S5B). Additionally, one of the top differentially expressed genes in the HSPC-like population compared to normal bone marrow HSPCs is *CD63* (Figure S5B). It encodes a cell surface protein that has been previously shown to distinguish normal and malignant hematopoietic cells (Mirkowska et al., 2013). Finally, the isolation of *KMT2A* fusion transcripts from this population further suggests that it is part of the leukemic population. The paradigm of leukemic stem cells (LSCs) was established largely based on study of AML and chronic myeloid leukemia (CML) (Vetrie et al., 2020). In contrast, LSCs in B-ALL is poorly characterized although the existence of LSCs is supported by the leukemia-initiating potential in xenotransplantation experiments (Castor et al., 2005; Elder et al., 2017; Rehe et al., 2013; le Viseur et al., 2008). Further experiments are needed to confirm the leukemia-initiating potential of the HSPC-like population identified in this study.

Given our observation of the large developmental spectrum of arrested blasts, the HSPC-like blasts are unlikely to be located at the apex of a hierarchy of LSCs in *KMT2A-r* leukemia. Instead, our data supports a model of a combination of transformed stem cells, multipotent and even unipotent progenitor cells that can support leukemogenesis and possibly mediate therapy resistance and immune evasion. However, several features of these HSPC-like blasts such as their canonical LSC/stem cell characteristics of quiescence, resistance to apoptosis and higher activity of ribosome biogenesis, suggest this population to be a drug resistant reservoir for relapse.

In addition to mediating chemotherapy resistance, the HSPC-like population may also play a role in immune evasion. A surprising recent discovery is that AML leukemic stem cells can actively evade immune cell killing by down-regulating surface expression of *NKG2D-L*, an activating signal for cytotoxic NK cells (Paczulla et al., 2019). Here, we report that an immunosuppressive signaling circuit in *KMT2A-r* leukemias, involving the HSPC-like blasts and cytotoxic lymphocytes NK cells and to a lesser extent CD8+ T cells. By leveraging single-cell data, we identified multiple ligand-receptor pairs between HPSC-like blasts and cytotoxic lymphocytes. By summing both activating and suppressive signaling L-R pairs, we found an overall suppressive signaling effect from HSPC-like blasts towards both cytotoxic lymphocyte types. An example of such suppressive signaling is the IFNG signaling between HSPC-like blasts and NK cells. We further identified an abnormal high level of IFNG in younger patients that is produced by the elevated activity of NK T cells in these patients. These results suggest modulation of interferon signaling and/or immune synapses as a new avenue of research for developing novel immunotherapeutic strategies for combating therapy-resistant *KMT2A-r* leukemia.

In summary, we identified higher lineage plasticity, specific resistance pathways and an age-specific immune-cell crosstalk in *KMT2A-r* ALL in very young infants. Across ages, we also found that epigenomic profiling more faithfully captures the degree of lineage plasticity (i.e. higher percentage of myeloid-like blasts in scATAC-Seq data, Figure 6A, 6H; Figure S7F). This is particularly striking in the cases of two patients who underwent lymphoid-to-myeloid lineage switch that was heralded by enhanced epigenetic plasticity months before the phenotypic event (Figure 6H, Figure S7F). Our findings have critical clinical implications as they support approaches geared towards targeted inhibitors or multi-target immunotherapy approaches rather than optimization of conventional stem cell-sparing chemotherapy or lineage-restricted immunotherapy, both of which fail to address the key underlying biological features of infant ALL.

## Acknowledgements

We acknowledge the Children’s Oncology Group and the Children’s Hospital of Philadelphia Center for Childhood Cancer Research biorepositories for provision of primary patient leukemia specimens. We thank Danika Makowski of the Center for Pediatric Tumor Cell Atlas for administrative support.

This work was supported by the National Cancer Institute (NCI) Human Tumor Atlas Network (HTAN) under award U2C CA233285 (to KT and SH). Additional support include NCI award CA243072 (to KT), grants from the Emerson Collective and the Coco Foundation (to KB), a Department of Defense CDMRP award W81XWH2010357 (to DB, EG, and PB), a National Human Genome Research Institute award HG010646 and a National Heart, Lung, and Blood Institute award DP2-HL142044 (to HW).

SPH is the Jeffrey E. Perelman Distinguished Chair in Pediatrics at The Children’s Hospital of Philadelphia.

## Author Contributions

KT, KB, and DB conceived the study. CC designed and performed experiments, analyzed and interpreted data, and wrote the manuscript. WY analyzed and interpreted data, and wrote the manuscript. YU, LF, YH, QZ developed analytic pipelines and performed data analysis. FA, QQ, CC, JF, PG, CL performed additional experiments. AF and IR performed data curation. PB, EG provided patient samples and interpreted data. KT, KB, and DB supervised the overall study, designed experiments, interpreted data. KT and KB wrote the manuscript. All authors approved the final version of the manuscript.

## Supplemental Titles and Legends

**Figure S1. Patient clinic data and quality assessment of single-cell omics data (related to Figure 1).**

**A)** Summary of patient clinical data, including leukemia subtype, fusion partner, age, and gender.

**B)** Representative FACS plot for sorting of whole bone marrow hematopoietic compartment (CD45+), lineage- CD34+ progenitors, and lineage-, CD34+, CD38-stem cell enriched population.

**C)** Summary of scRNA-Seq data after QC and removal of low-quality cells. Top, number of cells sequenced per sample; Bottom, number of genes identified per cell. Red horizontal line, average value across all samples.

**D)** Summary of scATAC-Seq data after QC and removal of low-quality cells. Top, number of cells sequenced per sample; Middle, number of uniquely mapped fragments per cell; Bottom, Fraction of accessible fragment reads located in ATAC-Seq peaks (FRIP). Red horizontal line, average value across all samples.

**E)** UMAP of scRNA-Seq and scATAC-Seq data from the infant cohort. Left panel, scRNA-Seq data; Right panel, scATAC-Seq data. Cells are colored by patient IDs. Total number of sequenced cells are indicated. *KMT2A* fusion partners are indicated in the parenthesis after patient IDs.

**F)** QC metrics of snmC-Seq2 data. Top left, number of total reads sequenced per cell by sample; top right, percentage of uniquely aligned reads per cell by sample; bottom left, percentage of genome coverage per cell by sample; bottom right, percentage of methylated CpG sites per cell by sample. Red horizontal lines, average values across all samples.

**Figure S2. Analysis of inter-tumor heterogeneity of infant** *KMT2A-r* **leukemia patient cohort (related to Figure 1).**

**A)** Genes ranked by standard deviation of mean expression across 18 infant samples. Top ranked B cell developmental and leukemia-related genes are highlighted in red.

**B)** Enriched pathways (FDR < 0.1) among top 50 genes ranked in panel A.

**C)** Violin plots for expression of top 25 genes ranked in panel A in CD19+ blasts from the 18 infant samples. Red vertical dashed line indicates division of the two age groups.

**D)** Aggregated scATAC-Seq signals at the *LYZ* locus in 18 infant samples. A transcriptional enhancer with varied activity across patients is highlighted in the black box.

**Figure S3. Normal hematopoietic trajectory and projection evaluation (related to Figure 2).**

**A)** Genome browser tracks for aggregated ATAC-Seq signals at representative cell-type-specific marker genes in healthy donors.

**B)** Motif accessibility enrichment computed by chromVAR for known lineage specific TFs, including myeloid (CEBPA), B lineage (EBF1), T lineage (EOMES), and hematopoietic stem and progenitor cells (GATA1).

**C)** Scatter plot for fraction of CD19+ blasts projected to cell types in the normal hematopoiesis developmental trajectory. X-axis, projected fraction in scRNA-Seq data. Y-axis, projected fraction in scATAC-Seq data.

**Figure S4. Transcriptomic and epigenomic signatures of blasts in younger patients (related to Figure 3).**

**A)** Enriched pathways among DEGs in additional clusters in Figure 3A.

**B)** Heatmap of differentially methylated regions (DMRs) overlapping with promoters of DEGs in blasts across B-cell developmental stages between younger and older patients. DMRs were identified using MethPipe (p-value < 0.05 & total CpG sites > 4 & differential CpG sites > 2). Red, up-regulated DEGs overlapped with hypomethylated DMRs in younger patients; Blue, down-regulated DEGs with hypermethylated DMRs in younger patients.

**C)** Example DEGs that overlapped with DMRs in pre-pro-B-like blasts. Top, *NR3C1*; Middle, *KLF9*; Bottom, *RUNX2*. Left panels, aggregated scATAC-Seq signals and methylation rates of blasts in younger and older patients. Right panels, normalized scRNA-Seq expression values of blasts in younger and older patients. Promoters are highlighted in yellow.

**D)** Methylation entropy difference of CpG sites in blasts between younger and older patients across B-cell developmental stages. All filtered CpG sites in the genome (STAR Methods) were used in the plot. P-values were based on *t*-test.

**Figure S5. A rare progenitor-like blast population in younger patients (related to Figure 4).**

**A)** Genome browser tracks for aggregated scATAC-Seq signals at representative HSPC-specific (*GATA2*, *CD34*, *HMGA2*) and B lineage-specific (*PAX5*, *VPREB1*, *DNTT*) marker genes.

**B)** Heatmap of differentially expressed genes between HSPC-like blasts and normal HSPCs using the cutoff of abs(Log2(FC) > 0.5 and FDR < 0.05. Values in the heatmap are row-wise Z-scores.

**Figure S6. Normal immune cell types in 18***KMT2A-r* **patients (related to Figure 5).**

**A)** UMAP of scATAC-Seq data for normal immune cells. Number of sequenced cells is indicated.

**B)** Frequency of normal immune cell types in younger and older patients.

**Figure S7. Characterization of longitudinal samples of two patients who had lineage switch after relapse and immunotherapy (related to Figure 6).**

**A)** Genome browser tracks for aggregated scATAC-Seq signals at representative B lineage (*CD19*, *VPREB1*, *CD79A*) and myeloid lineage-specific (*CD33*, *LYZ*, *CEBPA*) marker genes.

**B)** FACS plots showing CD19 and CD33 expression in bone marrow blasts of patient 1 before (top, sample ID 2184) and after lineage switch (bottom, sample ID 2263).

**C)** FACS plots showing CD19 expression in patient 2 at four time points. Top left, initial diagnosis ALL (blasts CD19+ on clinical flow, sample ID 1979); Top right, first relapsed ALL after chemotherapy and CD19 directed immunotherapy (blasts CD19 negative on clinical flow, sample ID 2524); Bottom left, second relapsed ALL (blasts CD19 negative on clinical flow, sample ID 2578); Bottom right, lineage switch to AML after CD22 targeted immunotherapy (AML, sample ID 2741).

**D)** & **E)** UMAPs of scRNA-Seq (D) and scATAC-Seq (E) data for a pediatric ALL *KMT2A-r* patient with leukemic samples at four time points, initial diagnosis, relapse 1 post chemotherapy and CAR19 therapy, relapse 2 post inotuzumab therapy, and after lineage switch to AML post CAR22 therapy. Left panel, UMAP of longitudinal samples, colored by assigned cell types. Total numbers of sequenced cells are indicated. Right panel, projection of patient cells to the normal hematopoietic trajectory. Grey dots, cells from healthy donors; colored dots, patient cells.

**F)** Fraction of B-, myeloid-lineage, and LMPP blasts at four time points. Top panel, fraction based on scRNA-Seq data; Bottom panel, fraction based on scATAC-Seq data.

**G)** Violin plots for representative signature genes in blasts from the four time points.

**Figure S8. LMPP-like blasts in *KMT2A-r* leukemia patients (related to Figure 6).**

**A)** Fraction of LMPP-like blasts in *KMT2A-r* leukemia samples based on scRNA-Seq data.

**B)** Fraction of LMPP-like blasts in *KMT2A-r* leukemia samples based on scATAC-Seq data.

**C)** Scatter plot for frequencies of M-lineage blasts and LMPP-like blasts in *KMT2A-r* leukemia samples. Spearman correlation is shown.

**D)** Enriched pathways among differentially expressed genes between normal LMPPs from healthy donors and LMPP-like blasts in patient samples post lineage switch.

**Figure S9. List of differentially expressed genes between CD19+ blasts and normal cells of the same developmental stage (related to Discussion section).**

**A)** 148 DEGs that are shared across at least 4 B-cell developmental stages.

**B)** Shared DEGs are enriched for known drug targets.

**Table S1.** Patient and healthy donor clinical and biospecimen information.

**Table S2.** Marker genes used for annotating cell types in the single-cell data.

**Table S3.** List of antibodies used in this study.

**Table S4.** List of primers used in this study.

**Table S5.** List of differentially expressed genes (abs(log_2_FC) > 0.5 & FDR < 0.05) of blasts arrested across B-cell developmental stages between younger (< 6 months old) and older patients (> 6 months old). DEGs are organized into clusters identified by hierarchical clustering.

**Table S6.** List of TF motifs with differential accessibility in blasts arrested across B-cell developmental stages between younger and older patients.

**Table S7.** List of differentially methylated regions in blasts across B-cell developmental stages between younger and older patients.

**Table S8.** Full transcriptional regulatory network for HSPC-like blasts vs normal HSPCs comparison.

## STAR Methods

### Human biospecimens

Peripheral blood or bone marrow samples from patients with infant ALL enrolled on the Children’s Oncology Group) COG clinical trial AALL15P1 or banked at the Children’s Hospital of Philadelphia Childhood Cancer Research (CCCR) Registry and Biorepository were obtained with parent informed consent according to the Declaration of Helsinki and institutional review board approval from all participating centers. Mononuclear cells from patients enrolled on the CCCR protocol were prepared using a ficoll gradient and stored frozen in 10% DMSO. Patient and healthy donor sample information is listed in Table S1.

### Cell sorting

Cells were thawed and stained with appropriate antibodies and DAPI (Cat #: 1246530100, Sigma) and immediately subjected to FACS sorting (FACSAria Fusion, BD). Sorted CD19+ and CD19-populations were subjected to scRNA-Seq and scATAC-Seq. For snmC-Seq2 protocol, sorted CD19+ were used. For healthy donor samples, DAPI-, DAPI-Lin-CD34+ and DAPI-Lin-CD34+CD38-populations were sorted and subjected to scRNA-Seq and scATAC-Seq. Antibody information is listed in Table S3.

### scRNA-Seq

Sorted cells were immediately processed using the 10x Genomics Chromium controller and the Chromium Single Cell 3’ Reagent Kits V3 protocol. 7,000-16,000 cells were loaded for each sample. Cells were partitioned into gel beads, lyzed, and barcoded through reverse transcription. cDNA were purified and amplified using appropriate cycle number following the manufacturer’s protocol. Libraries were constructed using 10x Genomics Library Prep Kit. Library quality was checked using Agilent High Sensitivity DNA Kit and Bioanalyzer 2100. Libraries were quantified using dsDNA High-Sensitivity (HS) Assay Kit (Invitrogen) on Qubit fluorometer and the qPCR-based KAPA quantification kit. Libraries were sequenced on an Illumina Nova-Seq 6000 with 28:8:0:87 paired-end format.

### scATAC-Seq

Sorted cells were centrifuged at 300g for 5 min at 4°C. 45uL of chilled lysis buffer was added to cell pellets and mixed by pipetting gently three times, and incubated 3 min on ice. After incubation, 50uL of pre-chilled wash buffer was added without mixing and centrifuged immediately at 300g for 5 min at 4°C. 95uL supernatant was carefully discarded and 45uL pre-chilled diluted nuclei buffer (10x Genomics) was added without mixing and sample was centrifuged at 300g for 5 min at 4°C. The nuclei pellet was then resuspended in 7uL pre-chilled diluted nuclei buffer and nuclei concentration was determined using a Countess II cell counter (Invitrogen). 7,000-16,000 nuclei were used for the transposition reaction in bulk, and then loaded to the 10x Genomics Chromium controller. Cells were partitioned into gel beads, lyzed, and barcoded followed by the Chromium Single Cell ATAC Reagent Kits protocol. The barcoded and transposed DNA was amplified with appropriate sample index. Library quality was checked using Agilent High Sensitivity DNA Kit and Bioanalyzer 2100. Libraries were quantified using dsDNA High-Sensitivity (HS) Assay Kit (Invitrogen) on Qubit fluorometer and the qPCR-based KAPA quantification kit. Libraries were sequenced on an Illumina Nova-Seq 6000 with 49:8:16:49 paired-end format.

### snmC-Seq2

The plates were prepared in an AirCleanⓇ600 PCR Workstation to minimize environmental DNA contamination. 4 μL lysis mix (containing 2 μL M-digestion buffer, 0.2 μL Proteinase K (20 mg/mL, from Zymo kit, cat. #D5023), 1.8 μL diluted lambda DNA (0.01 pg/well)) was added to each well of 96-well plates before covering by sealing film (Thermo Scientific, cat. #ab-0558). Single DAPI- CD19+ cell was sorted into each well. After cell sorting, the plates were covered by sealing film and incubated at 50°C for 20 min (PK-digestion). All steps of library preparation were performed in an AirCleanⓇ600 PCR Workstation to minimize environmental DNA contamination. The snmC-seq2 library preparation was performed as previously described with minor modifications (Luo et al., 2018). Briefly, bisulfite conversion of single cells was carried out using Zymo EZ-96 DNA Methylation-Direct™ Kit (Deep Well Format, cat. #D5023). The DNA was eluted in 9 μL Low EDTA TE buffer (Swift Biosciences) plus 1 μL of an assigned random primer (final concentration 0.5 μM) (Table S4) to allow for later multiplexing (8-plex) reactions.

Before random-priming, plates were denatured by incubating at 95°C for 3min and were immediately chilled on ice for 2min. 10 μL enzyme mix was added to each well. The following program was used for random-priming: 4°C for 5min, ramp up to 25°C at 0.1°C/sec, 25°C for 5min, ramp up to 37°C at 0.1°C/sec, 37°C for 60min. Then 2 μL Exonuclease I (20 U/μL, NEB cat. # M0293L) and 1 μL Shrimp Alkaline Phosphatase (rSAP) (1 U/μL, NEB cat. # M0371L) was added to each reaction and followed by incubation in a thermocycler at 37°C for 30min. The pooled samples were purified once with 0.8x SPRI beads and eluted with 10 μL Low EDTA TE buffer. Eluted samples were transferred to new PCR strips for downstream steps.

Before Swift Adaptase reaction, samples were denatured in a thermocycler at 95°C for 3min and subsequently chilled on ice for 2min. Then 10.5 μL Adaptase master mix was added to each reaction. Reactions were incubated in a thermocycler at 37°C for 30min. Subsequently, 30 μL PCR mix was added to each well, followed by mixing with pipetting. The following program was used for indexing-PCR: 95°C for 2min, 98°C for 30sec, 15 cycles of (98°C for 15sec, 64°C for 30sec, 72°C for 2min), 72°C for 5min, and hold at 4°C. PCR products were cleaned with two rounds of 0.8x SPRI beads. Library quality was then checked using Agilent High Sensitivity DNA Kit and Bioanalyzer 2100. Libraries were quantified using dsDNA High-Sensitivity (HS) Assay Kit (Invitrogen) on Qubit fluorometer and the qPCR-based KAPA quantification kit. Libraries were sequenced on an Illumina Nova-Seq 6000 with 150:8:8:150 paired-end format.

### Single-cell targeted long-read sequencing with Oxford Nanopore technology

Barcoded full-length cDNA library from 10x Genomics protocol was amplified using the primer that targets *KMT2A* exon 7 (Table S4) and KAPA HiFi HotStart ReadyMix (KAPA, Cat: KK2600). The following program was used for PCR: 95°C for 3min, 12 cycles of (98°C for 20sec, 60°C for 15sec, 72°C for 5min), 72°C for 5min, and hold at 4°C. Samples were cleaned with Dynabeads Kilobase BINDER (Thermo Fisher, Cat: 6010) to enriched for biotinylated amplified cDNA. After Dynabeads cleanup, samples were appended with Illumina P5 and P7 adapter sequences using additional PCR reaction: 95°C for 3min, 15 cycles of (98°C for 20sec, 60°C for 15sec, 72°C for 5min), 72°C for 5min, and hold at 4°C. PCR products were cleaned with one round of 0.7x SPRI beads. PCR products were quantified using dsDNA High-Sensitivity (HS) Assay Kit (Invitrogen) on Qubit fluorometer. Nanopore libraries were prepared using Nanopore Ligation Sequencing Kit (Nanopore, Cat #: SQK-LSK109). Samples were sequenced on Nanopore GridION.

### scRNA-Seq data processing

scRNA-Seq data for each patient were first processed using cellranger (v3.1.0) followed by additional processing using customized R scripts. Specifically, we filtered out cells with fewer than 1500 UMIs or greater than 50000 UMIs. Cells with more than 15% of UMI from mitochondrial genes were removed for downstream analyses. We further removed red blood cells with UMI for *HBB* gene greater than or equal to 3. DoubletFinder(McGinnis et al., 2019) was used to remove cell doublets with 5% of expected rate of doublets. The filtered gene-by-cell count matrix was used for downstream analyses.

### Integrated analysis of scRNA-Seq from 18 infant *KMT2A-r* patients

The 18 filtered gene-by-cell count matrices were combined into a bigger gene-by-cell count matrix and converted to a Seurat object, with the fraction of mitochondrial UMI and sample information added as the metadata. We computed cell cycle S phase and G/M2 phase scores using the CellCycleScoring function in Seurat, with cell cycle genes downloaded from https://satijalab.org/seurat/v3.2/cell_cycle_vignette.html. We additionally computed a heat shock gene signature score using the AddModuleScore function in Seurat using genes associated with heat shock proteins from HGNC (https://www.genenames.org/data/genegroup/#!/group/582). Next, we normalized the data using the *NormalizeData* function in Seurat. We selected the top 3,000 variably expressed genes (VEGs) using the *FindVariableFeatures* function in Seurat with the default setting. The VEGs were further scaled by the *ScaleData* function in Seurat. We also regressed out confounding factors including the fraction of mitochondrial UMIs, the total number of UMIs per cell, S phase score, G2/M phase score and heat shock score. We then performed principal component analysis (PCA) using the *RunPCA* function in Seurat with 50 principle components (PCs). The data were clustered using the *FindNeighous* and *FindClusters* functions, with the resolution=0.4. To reduce batch effect, we re-selected VEGs across the clusters. To do so, we summarized the count matrix into a gene-by-cluster count matrix and normalized it using the *cpm* function in the edgeR package with the parameter prior.count = 1. The top 3,000 VEGs across the normalized gene-by-cluster count matrix were used. Data were scaled, regressed out for confounding factors followed by PCA and clustering analyses. VEGs that were expressed in fewer than 1% of cells were removed from the analysis. This process of re-selecting VEGs was repeated once and the resulting top 50 PCs were used in Uniform Manifold Approximation and Projection (UMAP) using the *RunUMAP* function in Seurat for visualization with the default setting.

### Construction of normal hematopoietic development trajectory using scRNA-Seq data from healthy donors

The 27 filtered (9 donors each with 3 sorted samples) gene-by-cell count matrices were combined into a bigger gene-by-cell count matrix. To prevent sorted progenitor cells from dominating the cell composition, we down-sampled the sorted cells to 12,000, about half of the live cells. The combined matrix was converted to a Seurat object, with the fraction of mitochondrial UMI and sample information added as the metadata. We next applied the same procedure as for integrating patient scRNA-Seq data, except using top 1,500 VEGs, the first 20 PCs, and only regressing out confounding factors of the total UMI count per cell and the fraction of mitochondrial UMIs per cell.

### Cell type annotation for scRNA-Seq data

The cells of integrated patient data were annotated into Blasts, Monocytes, Mature B, T and NK cells based on marker gene expression. The normal hematopoietic trajectory built using healthy donor samples was further annotated into 17 cell types. The detailed cell types and corresponding marker genes used are summarized in Table S2.

### Projection of patient scRNA-Seq data onto the normal developmental trajectory

To project the patient cells onto the normal reference trajectory, we needed to transform the raw patient data as we did for constructing the normal developmental trajectory. Specifically, for each of the VEGs used for constructing the normal trajectory, we extracted the linear regression model and saved mean and standard deviation of the regression residuals. Then, for each corresponding gene in patient data, we subtracted the predicted score using the linear regression model and scaled the residuals with the corresponding mean and standard deviation. This transformed gene-by-cell matrix was then multiplied by the PCA feature loading matrix from normal trajectory to calculate the projected PC scores for each patient cell. Finally, the *umap_transfer* function from the uwot R package was used to generate the projected patient UMAP coordinates. The projected cell type of each patient cell is defined as the cell type of the closest (in Euclidean distance) healthy donor cell in the UMAP embedding.

### Identification of differentially expressed genes

For each comparison, we identified the differential expressed genes between two groups using the *FindMarkers* function in Seurat. We performed *LR* test on the raw counts and regressed out the total UMI count per cell as a confounding factor.

### scATAC-Seq data processing

scATAC-Seq data for each sample was first demultiplexed using the mkfastq function of cellranger-atac (v.1.1.0) tool. The fastq files were then processed using the scATAC-pro(Yu et al.) package with the default settings for most modules. Specifically, the cell barcode was written to the name of each read in the fastq files using the *demplx_fastq* module and then the reads were aligned to the hg38 genome assembly using the *mapping* module with bwa as the sequence aligner. Peaks were called using the *call_peak* module using the ‘COMBINED’ option. Peaks were called using MACS2 for each cell cluster. Peaks identified from different clusters were further merged if they were within 200bp of each other. With the called peaks, we then constructed the raw peak-by-barcode count matrix. Using the *call_cell* module, barcodes with more than 3,000 total fragments and the fraction of reads in peaks (FRiP) greater than 50% were identified as cells. The peak-by-cell count matrix was used for downstream analyses.

### Integrated analysis of scATAC-Seq from 18 infant *KMT2A-r* ALL patients

To integrate data from all patients, we first merged peaks called from all patients using the *mergePeaks* module of scATAC-pro. With this union set of peaks, we reconstructed the peak-by-cell count matrix for each patient using the *reConstMtx* module in scATAC-pro. The 18 peak-by-cell matrices were combined into a bigger peak-by-cell matrix and converted to a Seurat object. The data were next normalized using the TF-IDF normalization in scATAC-pro. Top 10,000 variably accessible peaks (VAPs) were identified using the *FindVariableFeatures* function in Seurat. We further filtered peaks that were accessible in fewer than 1% of the cells. TF-IDF normalization was then performed on the count matrix with only the VAPs. The PCA was performed on the VAPs and the cells were clustered using the first 50 PCs and the *FindClusters* function in Seurat with the resolution=0.4. To reduce batch effect, we reselect the VAPs across the clusters. To do so, we summarized the count matrix into a peak-by-cluster count matrix and normalized it using the *cpm* function in the edgeR package with parameter prior.count = 1. The top 10,000 VAPs across the normalized peak-by-cluster count matrix were selected and scaled, followed by PCA and clustering analyses. This process of re-selecting VAPs was repeated twice and the resulting first 50 PCs were used in Uniform Manifold Approximation and Projection (UMAP) using the *RunUMAP* function in Seurat for visualization with the default setting.

### Construction of normal hematopoietic development trajectory using scATAC-Seq data from healthy donors

Similar to integrating data from 18 patients, we first merged the peaks from all healthy donors using the *mergePeaks* module in scATAC-pro and reconstructed the peak-by-cell count matrix using combined peaks and the *reConstMtx* module in scATAC-pro. The 21 (7 healthy donors each with 3 sorted samples) peak-by-cell count matrices were then combined into a bigger peak-by-cell count matrix and converted to a Seurat object. We next applied the same procedure as integrating patient scATAC-Seq data, except for using the first 30 PCs.

### Projection of patient scATAC-Seq data onto the normal developmental trajectory

For each patient, we first reconstructed the peak-by-cell matrix using the VAPs used for constructing normal reference trajectory, using the *reConstMtx* module in scATAC-pro. The reconstructed matrix for each patient was normalized using the TF-IDF module in scATAC-pro. For each peak, we scaled the data using the mean and standard deviation calculated based on that peak in the healthy donor data. The transformed peak-by-cell matrix was then multiplied with the PCA feature loading matrix from normal reference to calculate the projected PC scores for each of the patient cells. Finally, the *umap_transfer* function from uwot R package was used to generate the projected patient UMAP coordinates. The projected cell type of each patient cell is defined as the cell type of the closest (in Euclidean distance) normal control cell in the UMAP embedding.

### Cell type annotation for scATAC-Seq data

The cells in the integrated patient data were annotated as Blasts, Monocytes, Mature B, T/NK cells based on TF enrichment score and gene activity score of marker genes. The TF enrichment score was computed using *chromVAR* and gene activity score was computed by summarizing fragments overlapped with the gene body or gene promoter (+/− 2kb around the TSS). The normal scATAC-Seq data was annotated for cell types by label transfer from normal scRNA-seq data using Seurat with the default setting.

### Identification of transcription factor motifs with differential genome-wide chromatin accessibility

We performed differential motif enrichment analysis based on the following comparisons: 1) between HSPC-like blasts and normal HSPCs; 2) between blasts arrested at a given development stage and corresponding normal cells at that stage; 3) between blasts of the same developmental stage in younger and older patients; 4) between blasts of the B-cell lineage and the myeloid lineage. For each cell group, we first used the chromVar algorithm to identify TF motifs that had enriched genome-wide chromatin accessibility in that cell group. We then tested differential enrichment between two cell groups using Wilcoxon test and the chromVar deviation scores. P-values were adjusted for multiple testing using the Benjamini-Hochberg method. Significant TF motifs were removed if the corresponding TF genes were expressed in fewer than 10% (20% for the comparison between HSPC-like blast and normal HSPC) of cells in the group.

### snmC-Seq2 data processing

Sequencing reads were demultiplexed using the 6 bp indices in the fastq files, followed by adapter trimming by cutadapt with the parameter settings: −f fastq −q 20 −u 10 −m 30 −a AGATCGGAAGAGCACACGTCTGAAC. Demultiplexed and trimmed reads were aligned using Bismark (Krueger and Andrews, 2011) with the default settings. Aligned reads were post-processed to remove ambiguous reads with mapping quality < 10, clonal reads, and reads containing three consecutive non-CpG methylation (indicating failure of bisulfite conversion). Aligned and filtered reads were counted using the countBam function and mapping rate was computed using the pileup function, both of which were implemented in the Rsamtools package (Morgan et al., 2016). For downstream analysis, we only used cells that had 200,000 filtered aligned reads and 20% mapping rate. We computed the methylation level of a region by calculating the ratio of methylated CpG sites to all methylation calls in that region. At least 10 overlapping CpG methylation calls were required to estimate the methylation levels of a region for each cell.

To integrate snmC-Seq and scATAC-Seq data, we first constructed a gene-by-cell matrix that consisted of the methylation level of the gene body and 2kb flanking region for each gene. For the missing value of a gene in a cell, we imputed it as the mean of observed methylation level of the rest of genes in the same cell. Next, we co-embedded snmC-Seq data with the scATAC-Seq data of blasts from the same 11 patients using Seurat. We first created a Seurat object that contained 1-log2(mtx+0.5), where the mtx was the imputed gene-by-cell methylation rate matrix. We then used *FindVariableFeatures, ScaleData and RunPCA* functions in Seurat to process the snmC-Seq data. For scATAC-Seq data, we constructed a gene activity-by-cell matrix using accessibility of gene body and 2kb flanking region and constructed an Seurat ‘assay’ called ACTIVITY in the original scATAC-Seq seurat object. This assay was subjected to normalization, scaling and dimensionality reduction using Seurat. The snmC seurat object was then co-embedded with the scATAC-Seq seurat object treating the scATAC-Seq data as the reference. A cell in snmC-Seq data was annotated based on its nearest cell in the scATAC-Seq data in the UMAP space.

For each developmental stage, we performed differential methylation analysis using MethPipe (Song et al., 2013) and aggregated cells from younger and older patients as the input. We then calculated the methylation rate for each CpG, differentially methylated CpGs (using p-value=0.05 as the cutoff) and differentially methylated regions (DMRs) using *methcounts*, *hmr, methdiff* and *dmr* functions in MethPipe, respectively. We further filtered DMRs to have at least 4 CpGs and at least 2 differentially methylated CpGs in the DMR. We computed the methylation entropy for each CpG using the *methentropy* function of MethPipe and the default sliding window size covering 4 CpGs. Because low sequencing coverage at a locus can cause inaccurate estimates of methylation entropy, we only calculated methylation entropy using CpGs having more than 100 reads in the sliding window.

### Analysis of Nanopore sequencing data

Oxford nanopore sequencing reads were demultiplexed using AmpBinner (https://github.com/WGLab/AmpBinner). AmpBinner first identified the p5 and read1 sequence of each read. The 20 bp sequence downstream of the read1 is expected to contain the barcode sequence and was extracted from each read and aligned to all possible cell barcodes in the sample. The alignment was performed using minimap2 (Li, 2018) with optimized parameters for short sequences (-k 3 -w 2 -n 1 -m 10 -s 40). The number of base edits (including mismatches, insertions, and deletions) were counted from the ‘CIGAR’ string. A read was confidently assigned to a cell barcode if the number of base edits was fewer than 3 and the second-best barcode had 3 or more base edits.

Demultiplexed sequencing reads were aligned to the human reference genome GRCh38 using minimap2 (Li, 2018) with parameters allowing detection of splicing isoforms (-ax splice -uf -k14). Fusion reads were called from the bam file using LongGF (https://github.com/WGLab/longgf) and AlignQC (Weirather et al., 2017). A read was identified as a fusion read if it was aligned to both the primer region and the gene body of the expected fusion partner. The read was identified as wild-type read if it was aligned to both the primer region and exon 15 of the *KMT2A* gene. For downstream analysis, only reads consistently called by both methods were considered fusion or wild-type reads.

### Prediction of enhancer-promoter interactions

Enhancer-Promoter (EP) interactions were predicted using a regression-based method described in (Zhu et al., 2020)A linear regression was conducted for each gene, with the gene expression in each cell as the dependent variable, and the normalized accessibility of the peaks within +/− 500kb of the gene promoter as the independent variables. If the regression coefficient of a peak is greater than 0.1 and the Benjamini-Hochberg adjusted p-value is smaller than 0.05, the EP pair was called significant. Cell-type-specific EP pairs were called if the enhancer peak had higher chromatin accessible (log_2_FC > 1) in the given cell type than the compared cell type(s).

### Construction of transcriptional regulatory network

The transcriptional regulatory network was constructed using differentially expressed genes and enriched TFs in HSPC-like blast compared to normal HSPC. A gene was defined as a target of a TF if there was a predicted HSPC-like-blast specific enhancer-promoter interaction and a TF motif hit at the enhancer. The color of the gene node is proportion to the −log10(FDR) in the DEG analysis, and the size of the TF node is proportion to the −log10(FDR) in the TF enrichment analysis. The thickness of an edge is proportional to the −log10(FDR) of the linear regression coefficient in the EP prediction analysis.

### Analysis of ligand-receptor interaction in signaling pathways

We identified ligand-receptor (L-R) interactions between blasts and cytotoxic lymphocytes using CellphoneDB (Efremova et al., 2020) and Kumar’s method (Kumar et al., 2018b). Normalized gene-by-cell expression matrix (down-sampled to 30K cells) and cell type annotation were used as the input to both methods. L-R interactions predicted by CellphoneDB with a p-value < 0.01 were considered significant. For Kumar’s method, we used the top 15% of predictions for each cell-type pair as significant interactions. Interactions found by both methods were considered as final predictions. For downstream analysis of comparing HSPC-like blasts and CD19+ blasts, among significant L-R pairs, we only considered those in which the ligand/receptor gene was differentially expressed (Wilcoxon test, FDR < 0.05) in one of the two blast types.

### Pathway enrichment analysis

To compute the pathway enrichment score for each cell, we used a modified approach based on AUCell. Instead of using all genes for ranking, we used an approach called IFF selection (Zhu et al., 2020) to remove housekeeping genes and keep cell-type specific genes. We then performed stage-wise Student’s *t*-test on the enrichment score for comparison between healthy donor HSPCs and patient HSPC-like blasts. Enriched pathways were identified using q-value <= 0.01.

### GO enrichment analysis

We used Metascape (Zhou et al., 2019) to perform GO enrichment analysis for differentially expressed genes using the default setting.

